# Serial KinderMiner (SKiM) Discovers and Annotates Biomedical Knowledge Using Co-Occurrence and Transformer Models

**DOI:** 10.1101/2023.05.30.542911

**Authors:** Robert J. Millikin, Kalpana Raja, John Steill, Cannon Lock, Xuancheng Tu, Ian Ross, Lam C Tsoi, Finn Kuusisto, Zijian Ni, Miron Livny, Brian Bockelman, James Thomson, Ron Stewart

## Abstract

**Background:** The PubMed database contains more than 34 million articles; consequently, it is becoming increasingly difficult for a biomedical researcher to keep up-to-date with different knowledge domains. Computationally efficient and interpretable tools are needed to help researchers find and understand associations between biomedical concepts. The goal of literature-based discovery (LBD) is to connect concepts in isolated literature domains that would normally go undiscovered. This usually takes the form of an A-B-C relationship, where A and C terms are linked through a B term intermediate. Here we describe Serial KinderMiner (SKiM), an LBD algorithm for finding statistically significant links between an A term and one or more C terms through some B term intermediate(s). The development of SKiM is motivated by the the observation that there are only a few LBD tools that provide a functional web interface, and that the available tools are limited in one or more of the following ways: 1) they identify a relationship but not the type of relationship, 2) they do not allow the user to provide their own lists of B or C terms, hindering flexibility, 3) they do not allow for querying thousands of C terms (which is crucial if, for instance, the user wants to query connections between a disease and the thousands of available drugs), or 4) they are specific for a particular biomedical domain (such as cancer). We provide an open-source tool and web interface that improves on all of these issues.

**Results:** We demonstrate SKiM’s ability to discover useful A-B-C linkages in three control experiments: classic LBD discoveries, drug repurposing, and finding associations related to cancer. Furthermore, we supplement SKiM with a knowledge graph built with transformer machine-learning models to aid in interpreting the relationships between terms found by SKiM. Finally, we provide a simple and intuitive open-source web interface (https://skim.morgridge.org) with comprehensive lists of drugs, diseases, phenotypes, and symptoms so that anyone can easily perform SKiM searches.

**Conclusions:** SKiM is a simple algorithm that can perform LBD searches to discover relationships between arbitrary user-defined concepts. SKiM is generalized for any domain, can perform searches with many thousands of C term concepts, and moves beyond the simple identification of an existence of a relationship; many relationships are given relationship type labels from our knowledge graph.

## Background

The goal of literature-based discovery (LBD) is to identify novel relationships between drugs, diseases, genes, and other entities that have gone unnoticed in corpora of text (1, 2). Typically, these are A-B-C relationships, meaning there is a relationship between the A and B entities and a relationship between the B and C entities, thus implying an (indirect) relationship between the A and C entities. These indirect relationships are often useful discoveries, but can be difficult to find; researchers must be familiar with the body of literature in different fields to connect disparate knowledge. Since the first LBD techniques used by Swanson in 1986 (3, 4), the LBD field has grown in tandem with increases in computational resources. LBD typically utilizes co-occurrence, semantic, and graph-based methods, often involving machine-learning (ML) and natural language processing techniques (2, 5, 6).

Recently, we have described KinderMiner (KM) (7, 8), a simple co-occurrence modeling algorithm, which can query a corpus of text using any user-defined search terms to find relationships between A and B terms using Fisher’s Exact Test. We decided to develop an extension of KM that finds A-B-C relationships based on four observations of available LBD tools: 1) they identify a relationship but not the type of relationship, 2) they do not allow the user to provide their own lists of B or C terms, hindering flexibility, 3) they do not allow for querying thousands of C terms (which is crucial if, for instance, the user wants to query connections between a disease and the thousands of available drugs), or 4) they are specific for a particular biomedical domain (such as cancer). We endevoured to provide an open-source tool that improves on all of these issues. In this work, we describe Serial KinderMiner (SKiM), which can perform LBD by searching for statistically significant A-B-C relationships in a corpus of text, provided A, B, and C search terms. SKiM is an LBD system that allows both “open” and “closed” discovery strategies. In closed discovery, the system is given an A term and C term, and the goal is to find B terms that link these two concepts together (leading to mechanistic insights, for example). In open discovery, the system is given an A term, and then searches for A-B links through a list of B terms, and each of the found B terms are paired with each term in a list of C terms. The goal of open discovery is to find new unknown C terms linked to the A term through some B intermediate, and thus is designed for hypothesis generation.

While KM and SKiM can find links between concepts, these algorithms report *p*-values generated by Fisher’s Exact Test, which do not describe the nature of the relationships between concepts. To augment these reported *p*-values, we have constructed a large knowledge graph from ML-extracted knowledge from the entire PubMed abstract corpus (for a review of biomedical knowledge graphs see Kilicoglu *et al*. (9), and Nicholson *et al*. (10)). The entities and relationships stored in the knowledge graph were extracted by a fine-tuned large language model, a strategy being employed by others for a variety of corpora and tasks (10, 11, 12, 13, 14). Once SKiM has completed its co-occurrence modeling search, this knowledge graph is queried to provide qualitative labels for statistically significant relationships, which help users semi-automatically interpret their results. The results display PubMed identifiers (PMIDs) so that users may investigate these putative relationships manually.

While LBD has a long and rich history, the availability of functional web-based tools for performing LBD is very restricted (1). Additionally, many previously published LBD tools and techniques are limited by restricted vocabularies and difficult-to-use or nonexistent user interfaces. The SKiM algorithm is open-source, and its implementation as a Dockerized web application programming interface (API) server (written in the Python programming language) can be found at https://github.com/stewart-lab/fast_km. A website that provides a public, easy- to-use interface to search PubMed abstracts using the KM and SKiM algorithms can be found at https://skim.morgridge.org. Lists of genes, drugs, transcription factors, symptoms/phenotypes, and diseases are provided as templates from which to build queries that may be interesting to users. Users can search for any text string(s) they like. Links to PubMed are embedded in the web interface that displays PMIDs tied to search results and relationships, allowing easy validation and exploration. The website allows users to filter, search, sort, share, and download the resulting lists of associations.

Of the LBD Systems listed in Table 2 of Gopalakrishnan *et al*. (1), we find that only LION LBD (specific for cancer biology) (15) and BITOLA (16) are functional and provide open discovery functionality comparable to SKiM; Arrowsmith (17, 18) provides closed discovery functionality. We compare SKiM to the BITOLA, LION LBD, and Arrowsmith tools on one LBD task. While none of these comparisons are ideal, from these experiments with other LBD tools, we conclude that SKiM fills a missing gap in the LBD toolbox—namely: the user can easily perform open searches with large lists of user-defined B and C terms, and get relationship annotations from our knowledge graph.

## Implementation

### Serial KinderMiner (SKiM) algorithm

SKiM is a serial form of the KinderMiner co-occurrence based algorithm described previously (8). SKiM is intended to find relationships between A and C terms through B-term intermediates. This is accomplished via co-occurrence modeling. Users provide lists of A, B, and C terms as inputs; SKiM then pairs each A term with each B term and performs a Fisher’s Exact Test to compute a *p*-value for each pair. A-B pairs that meet a *p*-value threshold (1 ×10^−5^ by default, maximum of 300 A-B pairs) move on to the next stage, where those B terms are paired with all C terms. Fisher’s Exact Test is then performed for each B-C pair. Thus, each A-B pair has a *p*-value, and each B-C pair has a *p*-value, subject to some filtering parameters. If an A-B pair and a B-C pair both have low *p*-values, and these pairs share a B term, then the A and C terms are considered to be putatively related through the intermediate B term. Each A, B, and C term can be a word, a phrase, or a set of words. Results are ordered by a prediction score, which is based on a combination of the FET *p*-value and the ratio of (B+C PMID counts/C PMID counts) for the best B-C pair. A detailed description of the prediction score is provided in the **Supplemental Methods**. In all tables in this manuscript, if a *p*-value is displayed as 0, this indicates that the *p*-value is below 2.2×10^−308^.

### Building the PubMed abstract index

Files containing PubMed abstracts were downloaded from https://ftp.ncbi.nlm.nih.gov/pubmed/ in extensible markup language (XML) format. Each abstract (title and text) was lowercased and subsequently tokenized via the natural language toolkit (NLTK) Python package. Each token was saved to a lookup table (i.e., a dictionary) along with its PubMed identifier (PMID) and position of the token in the abstract. The lookup table containing all tokens from all PubMed abstracts was written to disk as a 74 gigabyte memory-mapped file for random access.

### SKiM web API server implementation

The SKiM algorithm was implemented in a web API server where incoming queries (“jobs”) are added to a work queue via hypertext transfer protocol (HTTP) request. Worker processes then retrieve jobs from the queue and perform the computations necessary to complete each job (lowercasing and tokenizing the A, B, and C terms, querying the PubMed abstract index, computing *p*-values via Fisher’s Exact Tests, and performing knowledge graph lookups). A job’s status, progress, and results can be retrieved via another HTTP request. Workers can perform other jobs besides SKiM jobs, such as KM jobs and building a new PubMed abstract text index. The server is intended to be run as a multicontainer Docker application; Python code and Docker build scripts can be found at https://github.com/stewart-lab/fast_km.

### Knowledge graph generation

A transformer ML model (19) was used for named entity recognition (NER) and relation extraction (RE). The PubMedBERT model from Microsoft (13) was fine-tuned on NER and RE tasks using spaCy. (20) For the NER task, 265 abstracts were hand-annotated using Prodigy (21) with 5 entity labels (“GGP” [gene or gene product], “CONDITION”, “CHEMICAL”, “BIO_PROCESS”, and “DRUG”). For the RE task, 2,573 sentences from 335 abstracts were annotated with 10 relationship labels (“ACTIVATES”, “INHIBITS”, “REGULATES”, “POS_ASSOCIATION”, “NEG_ASSOCIATION”, “COREF”, “BINDS”, “DRUG_INTERACTION_WITH”, “TREATS”, and “MUTATION_AFFECTS”). These labeled data were split by PMID into training (90%) and validation (10%) sets.

NER and RE spaCy models were trained on one Tesla T4 graphical processing unit (GPU) using Google Colab in ∼1 hour using default spaCy hyperparameters. These models were used to extract entities and relationships from 34 million PubMed abstracts, which took ∼72 hours on a distributed cluster of 32 GPU-enabled machines via the Center for High Throughput Computing at the University of Wisconsin–Madison.(22) Training data and config files are located at https://github.com/stewart-lab/kinderminer_kg. Extracted entities (376,386) and relations (3,953,657) were stored as a graph in a Neo4j database, where the named entities were stored as nodes and relations were stored as edges between nodes.

## Results and Discussion

### Evaluating SKiM on Classic Discoveries by Swanson and Smalheiser

For a given A term, open discovery SKiM uncovers associated C terms through intermediate B terms (A→B→C) (Figure 1). SKiM counts PubMed abstracts with the co-occurring A-B or B-C pairs and filters statistically significant pairs using the one-sided Fisher’s Exact Test (FET) *p*-value (see **Implementation**). In this manuscript, “significant” relationships are defined as relationships that achieve an FET *p*-value of less than 1×10^−5^.

**Fig. 1.**
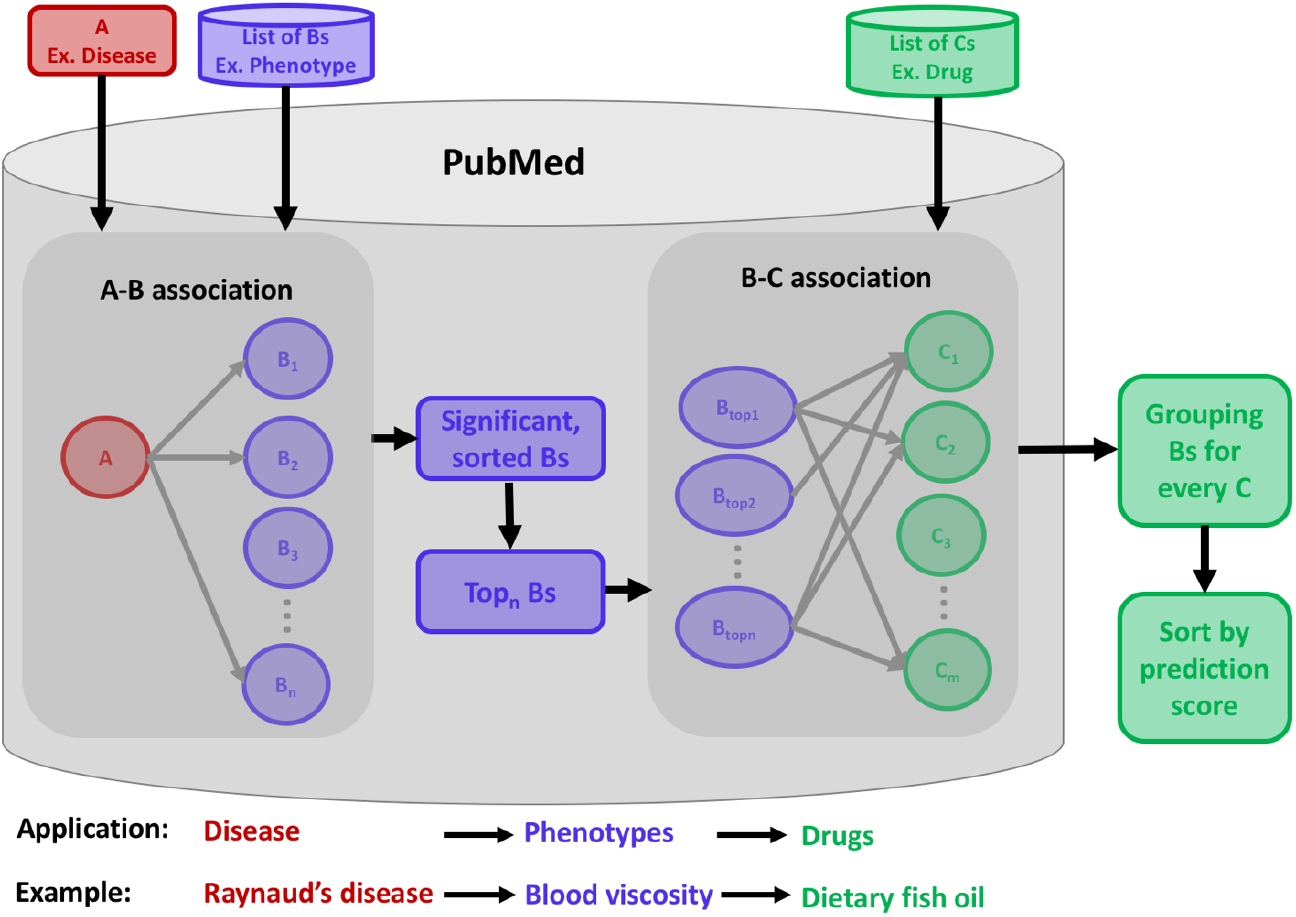
Visual depiction of the SKiM algorithm. An A term is provided by the user (for example, a disease such as “Raynaud’s”). A list of B-terms (e.g., phenotypes) are each paired with the A-term, and a Fisher’s Exact Test is performed to get a *p*-value. The significant A-B pairs are ranked and filtered by *p*-value and are each paired with a list of C terms (e.g., drugs). B-C pairs are sorted by prediction score. The user can provide their own B and C terms or can use pregenerated lists stored on the SKiM website.

We first evaluated SKiM by attempting to rediscover five relationships originally found in seminal work by Swanson and Smalheiser (4, 23, 24, 25, 26). So as to not bias the results, SKiM was only allowed to query abstracts that were published prior to each discovery. We used our phenotypes and symptoms list as the B terms, and a subset of 9,665 drugs annotated with diseases, genes, phenotypes, variants, or haplotypes from various expert-curated resources as C terms (see **Supplemental Methods** for details). Swanson and Smalheiser identified drugs, hormones, inflammation, and other concepts as B terms in their original discoveries; SKiM found all of these at a FET *p*-value less than 1×10^−5^ (Table 1).

**Table 1.** Rediscovery of five classic LBD findings by Swanson and Smalheiser, using SKiM. SKiM successfully rediscovers five of Swanson and Smalheiser’s discoveries when it is only allowed to search abstracts published prior to each of the original discoveries.

| A term | Cutoff date | Phenotype or Symptom (B) | A-B $p$-value | Swanson-Smalheiser Discovery (C) | B-C $p$-value | B-C Prediction Score Rank |
| --- | --- | --- | --- | --- | --- | --- |
| Raynaud’s | 1985 | Blood viscosity | $2 \times 10^{-33}$ | Dietary fish oil | $5 \times 10^{-10}$ | 462 |
| migraine | 1987 | Seizures | $6 \times 10^{-24}$ | Magnesium | $7 \times 10^{-36}$ | 822 |
| | | Depression | $2 \times 10^{-36}$ | | $2 \times 10^{-18}$ | |
| | | Muscle spasms | $7 \times 10^{-21}$ | | $3 \times 10^{-12}$ | |
| | | Tension | $6 \times 10^{-106}$ | | $1 \times 10^{-7}$ | |
| Alzheimer’s | 1995 | Normal gait | $5 \times 10^{-280}$ | Indomethacin | $5 \times 10^{-229}$ | 253 |
| | | Onset | $5 \times 10^{-128}$ | | $9 \times 10^{-38}$ | |
| | | Depression | $4 \times 10^{-96}$ | | $5 \times 10^{-14}$ | |
| | | Central nervous system disease | $3 \times 10^{-50}$ | | $4 \times 10^{-6}$ | |
| Alzheimer’s | 1995 | Normal gait | $5 \times 10^{-280}$ | Estrogen | 0 | 95 |
| | | Depression | $4 \times 10^{-96}$ | | 0 | |
| | | Onset | $5 \times 10^{-128}$ | | $9 \times 10^{-286}$ | |
| | | Dermal atrophy | $3 \times 10^{-127}$ | | $8 \times 10^{-43}$ | |
| | | Central nervous system disease | $3 \times 10^{-50}$ | | $3 \times 10^{-11}$ | |
| Somatomedin C | 1989 | Normal gait | $2 \times 10^{-192}$ | Arginine | 0 | 246 |
| | | Paraganglioma | $2 \times 10^{-33}$ | | 0 | |
| | | Glucose intolerance | $5 \times 10^{-65}$ | | 0 | |
| | | Cognitive abnormality | $4 \times 10^{-32}$ | | 0 | |

Importantly, SKiM also suggested some drug candidates that are likely to be false positives; thus, SKiM is intended as a mechanism to suggest reasonable candidates (i.e., a hypothesis generator) that can be validated by manual inspection of article texts, evaluation of information available in the SKiM knowledge graph, other sources of prior knowledge, and/or by wet-lab validation.

### Application of SKiM to Drug Repurposing

In the prior section, we focused on using open discovery SKiM to replicate five discoveries by Swanson and Smalheiser using a cutoff date of one year before each discovery. Here, we used open discovery SKiM to search for candidates for drug repurposing for four diseases from Swanson and Smalheiser’s work: Raynaud’s disease, migraine, Alzheimer’s disease, and schizophrenia. We assessed how far in advance SKiM found a disease–drug pair that later appeared in a clinical trial, limited to the top 100 drug hits for each disease.

In many cases, SKiM suggests drugs that had already been used to treat the disease or molecules otherwise associated with the disease (e.g., measurands); however, it does find drugs that were trialed up to 16 years after the cutoff date in the top 100 predictions. These predictions are shown in Table 2. In all cases, SKiM produces predictions before the first suggestion in the literature that the drug is a possible treatment for the disease, suggesting that SKiM will likely be useful for generating new hypotheses.

**Table 2.**
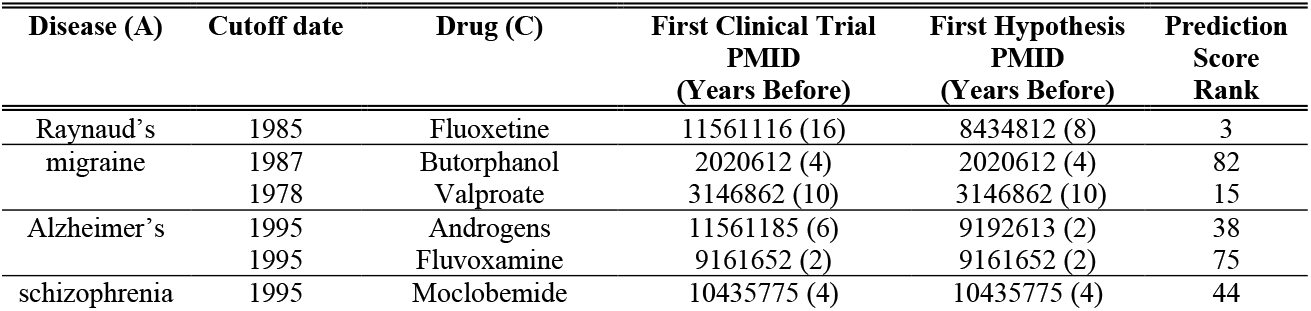
Significant disease–drug associations predicted by SKiM that were later tested in clinical trials. In all cases, the SKiM prediction is years before the hypothesis (that the listed drug might treat the disease) occurs in a PubMed abstract. Drugs in this table were limited to predictions that appeared in the top 100 hits ranked by prediction score.

### Application to Cancer Biology

SKiM is not limited to a specific application, such as drug repurposing; it accepts any combination of A, B, and C terms. To demonstrate SKiM’s generalizability, we used closed discovery SKiM to rediscover the LION LBD evaluation dataset related to cancer biology (15). With cutoff dates set to the year prior to discovery, SKiM found four out of five cancer biology discoveries from the LION LBD evaluation dataset; only one of the five discoveries was not significant (B-C *p*-value of 0.17) (Table 3; see **Supplemental Tables** for full query results and synonyms searched). Note, however, we are not able to recover this discovery (CXCL12→senescence←Thyroid cancer) using the closed discovery LION LBD interface either. This experiment demonstrates that SKiM is sensitive enough to find interesting and important A-B-C relationships in applications beyond drug repurposing.

**Table 3.** Rediscovery of LION LBD cancer biology discoveries using SKiM. Four of five discoveries are significant (A-B and B-C *p*-values < 1×10^−5^).

| A | Cutoff date | B | A-B $p$-value | C | B-C $p$-value |
| --- | --- | --- | --- | --- | --- |
| NF-κB | 2015 | Bcl-2 | $5 \times 10^{-76}$ | Adenoma | $1 \times 10^{-34}$ |
| NOTCH1 | 2015 | senescence | $6 \times 10^{-12}$ | C/EBPβ | $1 \times 10^{-14}$ |
| IL-17 | 2014 | p38α | $2 \times 10^{-8}$ | MKP-1 | $3 \times 10^{-9}$ |
| Nrf2 | 2010 | ROS | 0 | Pancreatic cancer | $5 \times 10^{-8}$ |
| CXCL12 | 2016 | senescence | $2 \times 10^{-7}$ | Thyroid cancer | 0.17 |

### Knowledge Graph Assists in Interpretation of Co-Occurrence Results

SKiM is a simple and sensitive text mining algorithm, as demonstrated earlier; however, it can lack interpretability in that it only reports *p*-values as its evidence for an association between two terms, and co-occurrence modeling cannot provide information on the nature of the relationship between the terms. We supplemented the SKiM co-occurrence modeling approach with entity (genes, drugs, biological processes, etc.) and relationship (treats, activates, inhibits, etc.) labels stored in a knowledge graph (Figure 2). These labels were extracted from PubMed abstract texts by natural language processing ML models (see **Implementation**). When a relationship is reported to the user, the PMIDs from which the relationship was extracted are displayed in the interest of transparency. The recall and precision of the named entity recognition (NER) and relationship extraction (RE) ML models are shown in Table 4.

**Table 4.** Precision, recall, and F1 score of the ML models used to build the knowledge graph. The PubMedBERT model from Microsoft was fine-tuned on NER and RE tasks using spaCy. The recall of the RE model is likely low because of sparse data, resulting in an unbalanced training set (i.e., most entities do not have a relationship with each other).

| Model | Precision | Recall | $F_1$ |
| --- | --- | --- | --- |
| NER | 0.885 | 0.814 | 0.848 |
| RE | 0.777 | 0.428 | 0.552 |

**Fig. 2.**
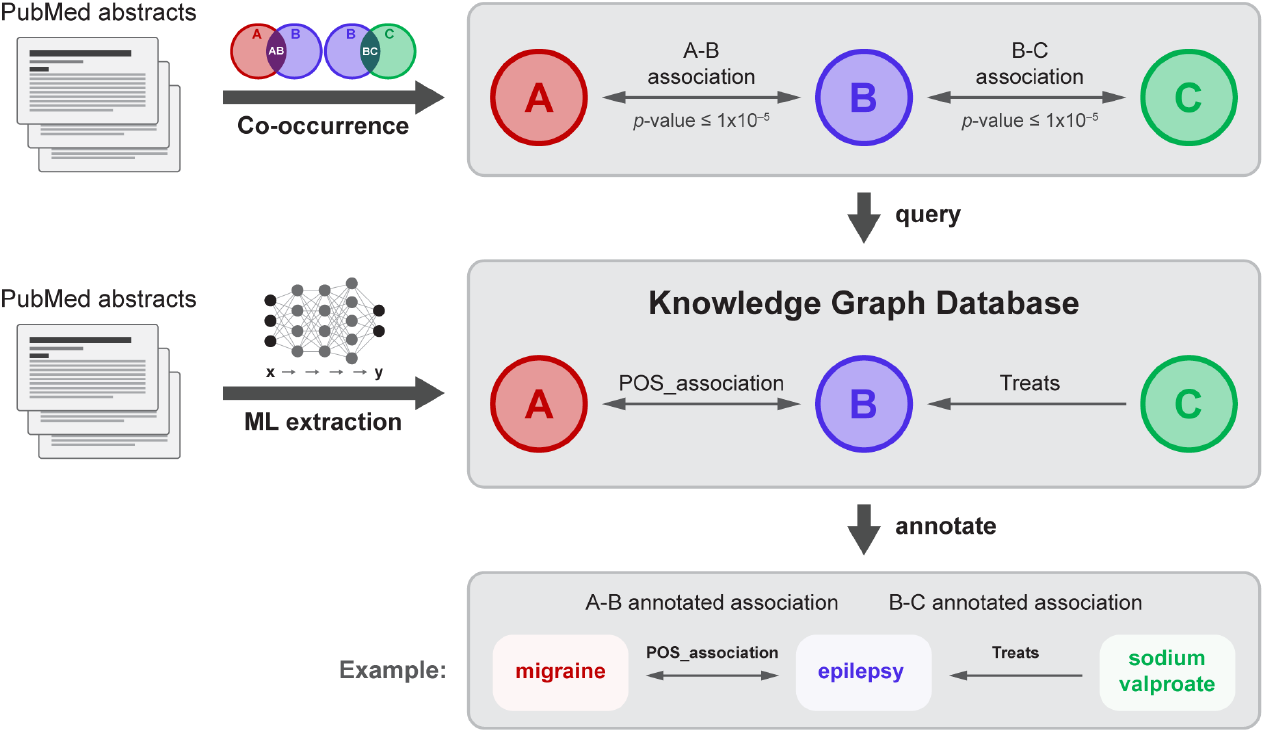
Visual depiction of SKiM coupled with knowledge graph annotation. Co-occurrence modeling (SKiM) is used to find statistically significant A-B-C relationships (e.g., migraine-epilepsy-sodium valproate). The knowledge graph, built by extracting biomedical entities and relationships from PubMed abstracts with ML, is queried for the A-B and B-C relationships. If these are found in the database, the relationships that SKiM found are annotated (e.g., “migraine POS_ASSOCIATION epilepsy,” “sodium valproate TREATS epilepsy”).

Note that the recall of the RE model is relatively low, though similar to recent work such as BioRED (27). (The low recall is likely due to the unbalanced/sparse nature of the training data; any two entities in a given sentence will likely not have a relationship). This means that only a minority of relationships that exist in PubMed are successfully extracted by the model. Even though only a subset of entities and relations are annotated in the knowledge graph, SKiM can still be used to find these significant relationships; they will simply lack annotations. Thus, we combine the sensitivity of simple text co-occurrence modeling and interpretability of natural language processing. We plan to expand the knowledge graph as better ML models become available.

As an example of the utility of the knowledge graph, SKiM (with a 1978 cutoff date) detects a statistically significant link between migraine and sodium valproate through the linking terms “seizure” and “epilepsy.” With only *p*-values provided, the user would need to conduct labor-intensive searches through several abstracts for each SKiM hit to discern the nature of each A-B and B-C relationship. Instead, these relationships are automatically labeled as “migraine (CONDITION) POS_ASSOCIATION epilepsy (CONDITION)” and “sodium valproate (DRUG) TREATS epilepsy (CONDITION)”; PMIDs are provided as evidence for these relationship labels. Similar labels are annotated for the migraine-seizure-sodium valproate relationships. It was not until 1988 that sodium valproate was used in a clinical trial to treat migraine (28).

An important feature of our knowledge graph is that the relationships are easily accessed via the SKiM web interface. Additionally, the interface allows users to access papers from which the relations of interest were extracted, a feature not available in most knowledge graphs (29). This latter feature is especially important because it makes the displayed relationships highly transparent to the user and allows manual validation.

### Comparison with Existing LBD Systems

We initially aimed to compare open-discovery SKiM with three existing LBD systems with functional web interfaces—BITOLA (16), LION LBD (15), and Arrowsmith (17, 18)—on the Swanson and Smalheiser discoveries. BITOLA was able to find either three or four out of five discoveries depending on the score that constitutes a discovery; however, BITOLA does not allow a cutoff date and thus can benefit from articles published after the discovery, thereby skewing results and preventing fair comparison with SKiM. Additionally, BITOLA is restricted to use only UMLS concepts as A, B, and C terms.

LION LBD is also restricted to certain entity types, allowing only annotated chemicals, diseases, mutations, genes, cancer hallmarks, and species as A, B, and C terms. The LION LBD web interface uncovered none of the five A-B-C links from the discoveries by Swanson and Smalheiser at the cutoff date of one year prior to the discovery, though this is likely because LION LBD is focused on cancer biology discoveries and can only display a small number of C terms in its graphical display. It should be noted that in a closed search mode, LION LBD displayed A-B-C relationships for four out of five discoveries; however, the scores of those relationships were zero or close to zero and it is not clear if these constitute a “hit.” Additionally, LION LBD does not seem to contain information from articles published after 2017.

Arrowsmith is a “closed” LBD system (A→B←C) but has a search mode (“one node search”) that can be used with short lists of C terms. These C terms can be selected from the MeSH tree or input manually. Our attempt to run Arrowsmith on a list of 9,665 drugs from our drugs lexicon terminated without success; however, Arrowsmith does successfully recover two out of five discoveries of Swanson and Smalheiser with smaller lists of C terms (we required a score of >0.1 to call a discovery). The “two-node” literature search of Arrowsmith takes both an A term and C term as input and reports linking B terms. While finding these B terms can lead to mechanistic insights, it cannot provide new drug candidates without the user executing a two-node search for every drug in a potentially very long list.

### SKiM Closed Discovery

SKiM provides both open discovery and closed discovery approaches to LBD. A closed search in SKiM can be performed by simply providing an A term, a C term (or short list of C terms), and then a list of B terms. As an example, the molecular mechanisms underlying the relationship between inflammation and cancer progression would not take shape in the literature until the 2000s (30). Using SKiM in a closed search mode, with “tumorigenesis” as the A term, “inflammation” as the C term, and genes as B terms, 14 genes were significantly associated with both tumorigenesis and inflammation when only searching abstracts published from 2001 and before. Of these 14 genes, 12 are true positives, with the remaining two discoveries being false positives because of semantic ambiguity (Table 5). Of the 12 true positive genes, six were not yet co-mentioned in an abstract with both tumorigenesis and inflammation in 2001. Of these six putative discoveries, five were later shown to link inflammation and cancer (31, 32, 33, 34, 35).

**Table 5.** Application of SKiM to a closed search problem. “Tumorigenesis” was the only A term searched, and “inflammation” was the only C-term searched; 17,545 genes were the B-terms. Only abstracts published in 2001 or before were searched. A total of six of the genes were already co-mentioned with both tumorigenesis and inflammation in the same article by 2001. Also, six were not co-mentioned with both tumorigenesis and inflammation in the same article by 2001; five of these latter six would later be shown to link tumorigenesis and inflammation. Finally, two genes (LOX and DES) were false positives caused by semantic ambiguity.

| A | Cutoff date | B | A-B *p*-value | C | B-C *p*-value | First A-C Publication (Year) |
| --- | --- | --- | --- | --- | --- | --- |
| Tumorigenesis | 2001 | AHR | $9 \times 10^{-7}$ | Inflammation | $6 \times 10^{-138}$ | 17848686 (2007) |
| | | CD44 | $6 \times 10^{-45}$ | | $5 \times 10^{-93}$ | 7530464 (1994) |
| | | FAS | $3 \times 10^{-23}$ | | $8 \times 10^{-107}$ | 10358186 (1999) |
| | | REL | $5 \times 10^{-10}$ | | $7 \times 10^{-35}$ | 15197457 (2004) |
| | | JUN | $4 \times 10^{-40}$ | | $1 \times 10^{-36}$ | 10657993 (2000) |
| | | FOS | $5 \times 10^{-42}$ | | $7 \times 10^{-38}$ | 10657993 (2000) |
| | | HGF | $5 \times 10^{-34}$ | | $9 \times 10^{-22}$ | 14596869 (2003) |
| | | STAT1 | $2 \times 10^{-7}$ | | $3 \times 10^{-13}$ | 16734720 (2006) |
| | | APC | $3 \times 10^{-263}$ | | $1 \times 10^{-26}$ | 8672984 (1995) |
| | | STAT3 | $4 \times 10^{-7}$ | | $4 \times 10^{-12}$ | 12219085 (2002) |
| | | LOX | $7 \times 10^{-6}$ | | $5 \times 10^{-9}$ | N/A |
| | | DES | $6 \times 10^{-28}$ | | $4 \times 10^{-23}$ | N/A |
| | | PCNA | $1 \times 10^{-59}$ | | $5 \times 10^{-12}$ | 20975039 (2010) |
| | | EGF | $5 \times 10^{-78}$ | | $1 \times 10^{-6}$ | 7895532 (1995) |

## Conclusions

SKiM is a simple but powerful and flexible open-source tool for finding known and novel associations between A and C terms, through B term intermediates. These terms can be concepts such as drugs, genes, diseases, or any word, phrase, or set of words. SKiM’s flexibility in searching for terms can be advantageous when a particular term is not included in a predefined list (“lexicon”). However, this approach does place the burden on the user to generate these terms if they are not already in a lexicon. Currently, we use the SKiM algorithm to search PubMed abstracts, though this corpus will likely expand in the future to other repositories of biomedical text such as Scopus. As we have demonstrated, SKiM can find relationships between entities without requiring that they conform to a predefined relationship type, using both open and closed search strategies. Thus, the co-occurrence algorithm employed by SKiM allows the user to cast a wide net to look for potential associations between entities. We supplement SKiM search results with qualitative entity and relationship labels extracted from PubMed abstracts with ML to aid in the interpretation of the co-occurrence-based results. The abstracts from which these entities and relationships were extracted are displayed on the SKiM website’s user interface for transparency.

In the future, we plan to make the knowledge graph of entities and relationship labels directly searchable and use it in other aspects of the SKiM website, such as generating predefined term lists for users to search. Additionally, the ML-generated labels could help users search within certain contexts; for example, “IF” is a difficult gene symbol to search when lowercased because it occurs in many contexts in which it is not used as a gene symbol. We plan to add a feature to SKiM in which a user could search only abstracts in which ambiguous terms like “IF” have been appropriately type-labeled by a ML model.

In summary, SKiM provides functionality that was previously missing from the LBD field. SKiM is generalized for any domain, can perform searches with many thousands of user-defined C term concepts, and moves beyond the simple identification of an existence of a relationship; many relationships between biomedical entities are given relationship type labels from our knowledge graph.

## Supporting information

Supplemental information for Table 3 and 5

## Availability and Requirements

**Project Name:** Serial KinderMiner (SKiM)

**Project home page:** https://skim.morgridge.org/ (user-facing website);

https://github.com/stewart-lab/fast_km (backend server code)

**Operating system(s):** Platform independent

**Programming language:** Python

**Other requirements:** To host a SKiM server, Python 3.8 or higher is required; to use the algorithm on the website, no requirements except a web browser

**License:** MIT

**Any restrictions to use by non-academics:** No restrictions

## List of Abbreviations

KM: KinderMiner
SKiM: Serial KinderMiner
LBD: literature-based discovery
FET: Fisher’s Exact Test
PMID: PubMed ID
XML: extensible markup language
NER: Named entity recognition
RE: Relation extraction
HTTP: Hypertext transfer protocol
NLTK: Natural language toolkit
GPU: Graphical processing unit
UMLS: Unified Medical Language System
NLM: National Library of Medicine
API: Application Programming Interface
ML: machine-learning

## Declarations

## Ethics approval and consent to participate

Not applicable

## Consent for publication

Not applicable

## Availability of data and materials

The search strings that support the findings of Table 1 and Table 2 of this study are available from the National Library of Medicine (NLM), but restrictions apply to the availability of these data, which were used under license for the current study, and so are not publicly available. Data are however available from the authors upon reasonable request and with permission of NLM. The search strings that support the findings of Table 3 and Table 5 of this study are included in this published article’s supplementary information files.

## Authors’ contributions

RM, KR, and RS designed the study and evaluation. RM, KR, RS, XT, and JS developed KinderMiner 2.0, FastKM, and SKiM. CL, JS, RS, XT, and RM built the web interface. RM, IR, and ML built versions of the local PubMed repository. RM built the knowledge graph. IR and KR contributed to integrate Allie with the PubMed repository for handling abbreviations. KR, LCT, and RS compiled the lexicons from various existing resources. KR, RS, ZN, FK, and LCT provided statistical support. BB provided support for computing resources and web hosting. JT provided biological inferences or interpretation of the results. RM, KR, and RS wrote the manuscript, and every author has reviewed the work.

## Acknowledgments

We gratefully acknowledge computing resources provided by the Center for High Throughput Computing at UW–Madison. We thank Amy Freitag and Alicia Williams for editorial assistance and Matt Stefely for assistance with Figure 2.

## References

1. Gopalakrishnan V, Jha K, Jin W, Zhang A. A survey on literature based discovery approaches in biomedical domain. J Biomed Inform. 2019;93:103141.

2. Thilakaratne M, Falkner K, Atapattu T. A Systematic Review on Literature-based Discovery. ACM Computing Surveys. 2019;52(6):1–34.

3. Smalheiser NR. Rediscovering Don Swanson: the Past, Present and Future of Literature-Based Discovery. J Data Inf Sci. 2017;2(4):43–64.

4. Swanson DR. Fish oil, Raynaud’s syndrome, and undiscovered public knowledge. Perspect Biol Med. 1986;30(1):7–18.

5. Lardos A, Aghaebrahimian A, Koroleva A, Sidorova J, Wolfram E, Anisimova M, Gil M. Computational Literature-based Discovery for Natural Products Research: Current State and Future Prospects. Front Bioinform. 2022;2:827207.

6. Zhao S, Su C, Lu Z, Wang F. Recent advances in biomedical literature mining. Brief Bioinform. 2021;22(3).

7. Kuusisto F, Ng D, Steill J, Ross I, Livny M, Thomson J, Page D, Stewart R. KinderMiner Web: a simple web tool for ranking pairwise associations in biomedical applications. F1000Res. 2020;9:832.

8. Kuusisto F, Steill J, Kuang Z, Thomson J, Page D, Stewart R. A Simple Text Mining Approach for Ranking Pairwise Associations in Biomedical Applications. AMIA Summits on Translational Science Proceedings. 2017;2017:166.

9. Kilicoglu H, Rosemblat G, Fiszman M, Shin D. Broad-coverage biomedical relation extraction with SemRep. BMC Bioinformatics. 2020;21(1):188.

10. Nicholson DN, Greene CS. Constructing knowledge graphs and their biomedical applications. Comput Struct Biotechnol J. 2020;18:1414–28.

11. Nicholson DN, Himmelstein DS, Greene CS. Expanding a database-derived biomedical knowledge graph via multi-relation extraction from biomedical abstracts. BioData Min. 2022;15(1):26.

12. Nadkarni R, Wadden D, Beltagy I, Smith N, Hajishirzi H, Hope T. Scientific language models for biomedical knowledge base completion: an empirical study. arXiv preprint. 2020(2106.09700).

13. Gu Y, Tinn R, Cheng H, Lucas M, Usuyama N, Liu X, Naumann T, Gao J, Poon H. Domain-Specific Language Model Pretraining for Biomedical Natural Language Processing. ACM Transactions on Computing for Healthcare. 2021;3(1):1–23.

14. Lee J, Yoon W, Kim S, Kim D, Kim S, So CH, Kang J. BioBERT: a pre-trained biomedical language representation model for biomedical text mining. Bioinformatics. 2020;36(4):1234–40.

15. Pyysalo S, Baker S, Ali I, Haselwimmer S, Shah T, Young A, Guo Y, Hogberg J, Stenius U, Narita M, Korhonen A. LION LBD: a literature-based discovery system for cancer biology. Bioinformatics. 2019;35(9):1553–61.

16. Hristovski D, Peterlin B, Mitchell JA, Humphrey SM. Using literature-based discovery to identify disease candidate genes. Int J Med Inform. 2005;74(2-4):289–98.

17. Swanson D, Smalheiser N. An interactive system for finding complementary literatures: a stimulus to scientific discovery. Artificial intelligence. 1997;91(2):183–203.

18. Smalheiser NR, Torvik VI, Zhou W. Arrowsmith two-node search interface: a tutorial on finding meaningful links between two disparate sets of articles in MEDLINE. Comput Methods Programs Biomed. 2009;94(2):190–7.

19. Vaswani A, Shazeer N, Parmar N, Uszkoreit J, Jones L, Gomez A, Kaiser Ł, Polosukhin I. Attention is all you need. Advances in neural information processing systems. 2017:30.

20. Honnibal M, Montani I, Van Landeghem S, Boyd A. spaCy: Industrial-strength Natural Language Processing in Python. 2020.

21. Montani I, Honnibal M. Prodigy: A modern and scriptable annotation tool for creating training data for machine learning models.

22. The Center for High Throughput Computing [Available from: https://doi.org/10.21231/GNT1-HW21.

23. Swanson DR. Migraine and magnesium: eleven neglected connections. 1988.

24. Smalheiser NR, Swanson DR. Indomethacin and Alzheimer’s disease. 1996.

25. Smalheiser NR, Swanson DR. Linking estrogen to Alzheimer’s disease: an informatics approach. 1996.

26. Swanson DR. Somatomedin C and arginine: implicit connections between mutually isolated literatures. Perspect Biol Med. 1990;33(2):157–86.

27. Luo L, Lai PT, Wei CH, Arighi CN, Lu Z. BioRED: a rich biomedical relation extraction dataset. Brief Bioinform. 2022;23(5).

28. Sorensen KV. Valproate: a new drug in migraine prophylaxis. Acta Neurol Scand. 1988;78(4):346–8.

29. Peng J, Xu D, Lee R, Xu S, Zhou Y, Wang K. Expediting knowledge acquisition by a web framework for Knowledge Graph Exploration and Visualization (KGEV): case studies on COVID-19 and Human Phenotype Ontology. BMC Med Inform Decis Mak. 2022;22(Suppl 2):147.

30. Coussens LM, Werb Z. Inflammation and cancer. Nature. 2002;420(6917):860–7.

31. Guarnieri T. Aryl Hydrocarbon Receptor Connects Inflammation to Breast Cancer. Int J Mol Sci. 2020;21(15).

32. Li X, Wang F, Xu X, Zhang J, Xu G. The Dual Role of STAT1 in Ovarian Cancer: Insight Into Molecular Mechanisms and Application Potentials. Front Cell Dev Biol. 2021;9:636595.

33. Lu R, Zhang YG, Sun J. STAT3 activation in infection and infection-associated cancer. Mol Cell Endocrinol. 2017;451:80–7.

34. Owusu BY, Galemmo R, Janetka J, Klampfer L. Hepatocyte Growth Factor, a Key Tumor-Promoting Factor in the Tumor Microenvironment. Cancers (Basel). 2017;9(4).

35. Zhao H, Wu L, Yan G, Chen Y, Zhou M, Wu Y, Li Y. Inflammation and tumor progression: signaling pathways and targeted intervention. Signal Transduct Target Ther. 2021;6(1):263.

