## Supplementary methods for "Serial KinderMiner (SKiM) Discovers and Annotates Biomedical Knowledge Using Co-Occurrence and Transformer Models"

**Title**

[10]Currently at Amazon

[‡] Authors contributed to this work equally

*Corresponding author

### Supplemental Methods

*Calculation of the prediction score:* The prediction score is based on combining the FET *p*-value with the ratio of (B-C counts/C-counts) as follows:

log_FET = (minimum(323, -log10(FET *p*-value))/323
(We cap the log_FET value at 323 to prevent numeric overflows. We then divide by 323 to assure that the log_FET value will range between 0-1.)

log_ratio calculation:
ratio = (B-C count/C count)

If ratio = 1:
    then log_ratio = 1
else:
    log_ratio = (minimum(323, -log10(1- ratio) × 2500))/323

(We multiply the -log10(1-ratio) by 2500 because analysis of median ratios indicates that multiplying by 2500 will typically bring this value into a range comparable to the -log10(FET p-value) and thus the FET and ratio will have similar contributions to the predictions score.)

Finally:
prediction_score = log_FET + log_ratio
(Prediction score will range between 0-2, with a maximum contribution of 1 from the FET and 1 from the ratio.)

### Lexicon Building

*Phenotypes and symptoms lexicon:* We compiled a phenotypes and symptoms lexicon from Human Phenotype Ontology (HPO) (1), Phenome Wide Association Studies (PheWAS) (2), and Online Mendelian Inheritance in Man (OMIM) (3). We downloaded the hp.obo file from http://human-phenotype-ontology.github.io/ and extracted 29,263 phenotypes and synonyms. We downloaded 414 phenotypes from OMIM (https://www.omim.org/) and 1,866 phenotypes from

PheWAS catalog (https://phewascatalog.org/phecodes). The identifiers in the HPO, OMIM, and PheWAS catalog are resource specific. We assigned a customized identifier and removed the duplicates across the resources. The process obtained 31,156 phenotypes and synonyms. None of the resources include blood viscosity from Swanson's discovery. HPO contains "blood hyperviscosity" and two synonyms, "hypercoagulability" and "thrombophilia." We extended the phenotypes and synonyms list by deriving new terms from the existing ones (i.e., "blood hyperviscosity" to "blood viscosity"). We used a list of 500 prefixes and suffixes related to the medical terminology (4, 5) to remove the prefix/suffix from phenotypes and synonyms. Our approach obtained 90,634 new terms. However, all the new terms are not phenotypes or symptoms (e.g., "plastic kidney" from "dysplastic kidney," "opathy" from "nephropathy," and "renal s" from "renal cysts"). We used KinderMiner to filter 3,314 new terms, mapping to at least two PubMed abstracts, and manually curated 1,771 new phenotypes that are not in HPO, OMIM, and PheWAS. We used the diseases lexicon to assign CUI to phenotypes and symptoms. All the phenotypes and symptoms do not map to CUI from the diseases lexicon. We assigned customized CUIs for those without a CUI. For every phenotype or symptom, we included all the synonyms from the diseases lexicon. The approach obtained a phenotypes and symptoms lexicon with 35,229 phenotypes, symptoms, and their synonyms (i.e., 9,272 phenotypes and symptoms).

*Drugs lexicon:* Our drugs lexicon (6) is from UMLS Metathesaurus 2018AB version (7), DrugBank 5.0 (8), and PharmGKB (9). It provides a comprehensive list of 693314 drugs and synonyms. In brief, we filtered the drugs and biologics from UMLS Metathesaurus based on four semantic types: clinical drug, antibiotic, pharmacologic substance, and immunological factor, and included the missing drugs and synonyms from DrugBank and PharmGKB. Including all the semantic types under Chemicals and Drugs will lead to many FPs because all chemicals are not therapeutic drugs (6). We assigned a customized drug ID instead of a CUI to accommodate drugs from DrugBank and PharmGKB that are not in UMLS Metathesaurus. The customized drug ID includes a prefix “CD” (e.g., CD00014279 for zotepine). Similar to the diseases lexicon, we combined the drugs and synonyms with multiple customized drug IDs and assigned new customized drug IDs with “CDM” prefix, where “M” stands for modified (e.g., CDM00000663 for metformin).

The drugs lexicon has a plethora of general drug classes (e.g., antihypertensive agents), general concepts (e.g., antigens), and pharmacologic substances that are not drugs (e.g., chemical buffer). For drug repurposing, we derived a subset of drugs associated with disease, gene, phenotype, variant, or haplotype from various resources. We retrieved 5,288 drugs associated with diseases from CTD (10), NDF-RT (11), DrugBank (8), and FDA (12). Many FDA-approved drugs are missing in CTD, NDF-RT, and DrugBank, and vice versa. We retrieved 7,293 drugs associated with genes from CTD, DrugBank, and PharmGKB (9). We retrieved 2,620 drugs associated with phenotypes from CTD. We retrieved 642 drugs associated with variants or haplotypes from PharmGKB. We included 3,080 annotated drugs from PharmGKB. We utilized the drugs lexicon to remove the overlapping duplicates across the resources by mapping the drugs to drug ID in the drugs lexicon. The process obtained 9,665 drugs.

*Diseases lexicon:* We compiled a diseases lexicon from UMLS Metathesaurus (7) and SNOMED CT (13). The 2018AB version of UMLS Metathesaurus was installed in Rich Release Format (RRF) using MetamorphoSys, a UMLS installation wizard. The resource contains 3.64 million health-related concepts (or 13.9 million unique concept names) from 201 source vocabularies (e.g., MeSH, RxNorm). Within UMLS Metathesaurus, we used MRCONSO, a resource on the concepts and MRSTY, a resource on the sematic types for the concepts. Though UMLS Metathesaurus includes a complete version of SNOMED CT, the default installation includes only 54 SNOMED concepts. SNOMED CT within UMLS Metathesaurus is installed by selecting a specific edition (i.e., SNOMED CT US Edition) during installation using MetamorphoSys. This version of SNOMED CT includes unique concept identifier (CUI) from UMLS Metathesaurus, and we assigned the semantic types to SNOMED CT concepts using CUIs.

We filtered the concepts belonging to semantic types under “Disorder”: UMLS Metathesaurus consists of 163,073 diseases (or 437,580 diseases and synonyms) and SNOMED CT consists of 146,436 diseases (or 470,076 diseases and synonyms). We combined the resources and removed the duplicates and 84 disease synonyms matching with English stop-words (e.g., in, present). The process obtained 273,710 diseases (or 640,386 diseases and synonyms). Of these, 7,450 diseases and synonyms include more than one CUI in UMLS Metathesaurus. For example, T2D and its synonyms are assigned with two CUIs, C0011860 and C4014362. We combined diseases and synonyms with more than one CUI and assigned a customized CUI which starts with “M” to represent modified “CUI.” All the semantic types from multiple CUIs are assigned to the modified CUI. Our disease lexicon consists of 264,983 diseases (or 640,386 diseases and synonyms).

### References


1. Kohler S, Carmody L, Vasilevsky N, Jacobsen JOB, Danis D, Gourdine JP, Gargano M, Harris NL, Matentzoglu N, McMurry JA, Osumi-Sutherland D, Cipriani V, Balhoff JP, Conlin T, Blau H, Baynam G, Palmer R, Gratian D, Dawkins H, Segal M, Jansen AC, Muaz A, Chang WH, Bergerson J, Laulederkind SJF, Yuksel Z, Beltran S, Freeman AF, Sergouniotis PI, Durkin D, Storm AL, Hanauer M, Brudno M, Bello SM, Sincan M, Rageth K, Wheeler MT, Oegema R, Lourghi H, Della Rocca MG, Thompson R, Castellanos F, Priest J, Cunningham-Rundles C, Hegde A, Lovering RC, Hajek C, Olry A, Notarangelo L, Similuk M, Zhang XA, Gomez-Andres D, Lochmuller H, Dollfus H, Rosenzweig S, Marwaha S, Rath A, Sullivan K, Smith C, Milner JD, Leroux D, Boerkoel CF, Klion A, Carter MC, Groza T, Smedley D, Haendel MA, Mungall C, Robinson PN. Expansion of the Human Phenotype Ontology (HPO) knowledge base and resources. Nucleic Acids Res. 2019;47(D1):D1018-D27.
2. Denny JC, Ritchie MD, Basford MA, Pulley JM, Bastarache L, Brown-Gentry K, Wang D, Masys DR, Roden DM, Crawford DC. PheWAS: demonstrating the feasibility of a phenome-wide scan to discover gene-disease associations. Bioinformatics (Oxford, England). 2010;26(9):1205-10.
3. Amberger JS, Bocchini CA, Schiettecatte F, Scott AF, Hamosh A. OMIM.org: Online Mendelian Inheritance in Man (OMIM(R)), an online catalog of human genes and genetic disorders. Nucleic Acids Res. 2015;43(Database issue):D789-98.
4. Stanfield PS, Hui YH, Cross N. Essential medical terminology. 4th Edition ed: Jones & Bartlett Learning; 2013.
5. Chabner D-E. The language of medicine. 11th Edition ed: Elsevier; 2014.
6. Raja K, Patrick M, Elder JT, Tsoi LC. Machine learning workflow to enhance predictions of Adverse Drug Reactions (ADRs) through drug-gene interactions: application to drugs for cutaneous diseases. Sci Rep. 2017;7(1):3690.
7. Bodenreider O. The Unified Medical Language System (UMLS): integrating biomedical terminology. Nucl Acids Res. 2004;32(Database issue):D267-D70.
8. Wishart DS, Feunang YD, Guo AC, Lo EJ, Marcu A, Grant JR, Sajed T, Johnson D, Li C, Sayeeda Z, Assempour N, Iynkkaran I, Liu Y, Maciejewski A, Gale N, Wilson A, Chin L, Cummings R, Le D, Pon A, Knox C, Wilson M. DrugBank 5.0: a major update to the DrugBank database for 2018. Nucleic Acids Res. 2018;46(D1):D1074-D82.
9. Barbarino JM, Whirl-Carrillo M, Altman RB, Klein TE. PharmGKB: A worldwide resource for pharmacogenomic information. Wiley Interdiscip Rev Syst Biol Med. 2018;10(4):e1417.
10. Davis AP, Grondin CJ, Johnson RJ, Sciaky D, McMorran R, Wiegers J, Wiegers TC, Mattingly CJ. The Comparative Toxicogenomics Database: update 2019. Nucleic acids research. 2019;47(D1):D948-D54.
11. Carter JS, Brown SH, Erlbaum MS, Gregg W, Elkin PL, Speroff T, Tuttle MS. Initializing the VA medication reference terminology using UMLS metathesaurus co-occurrences. Proceedings AMIA Symposium. 2002:116-20.
12. U.S. Food & Drug Administration 2019 [Drugs@FDA: FDA Approved Drug Products]. Available from: https://www.accessdata.fda.gov/scripts/cder/daf/.

13. De Silva TS, MacDonald D, Paterson G, Sikdar KC, Cochrane B. Systematized nomenclature of medicine clinical terms (SNOMED CT) to represent computed tomography procedures. Comput Methods Programs Biomed. 2011;101(3):324-9.
